# Evaluation of anticancer and anti-mitotic properties of quinazoline and quinazolino-benzothiadiazine derivatives

**DOI:** 10.1101/108654

**Authors:** Thoukhir B. Shaik, M. Shaheer Malik, Zaki S. Seddigid, Sunitha R Routhu, Ahmed Kamal

## Abstract

Cancer is one of the major health and social-economic problems despite considerable progress in its early diagnosis and treatment. Owing to the emergence and increase of multi drug resistance to various conventional drugs, and the continuing importance on health-care expenditure, many researchers have focused to develop novel and effective anticancer compounds. In the present study, a series of in-house synthesized quinazoline and quinazolino-benzothiadiazine derivatives were investigated for their anticancer efficacy against a panel of five cancer (DU145, MCF7, HepG2, SKOV3 and MDA-MB-231) and one normal (MRC5) cell lines. Among all the tested compounds, fifteen of them exhibited promising growth-inhibitory effect (0.15 - 5.0 μM) and induced cell cycle arrest in G2/M phase. In addition, the selected compounds inhibited the microtubule assembly; altered mitochondrial membrane potential and enhanced the levels of caspase-9 in MCF-7 cells. Furthermore, the active compound with combination of drugs showed synergistic effect at lower concentrations and the drug uptake was mediated through clathrin mediated endocytic pathway. Our results indicated that quinazoline and quinazolino-benzothiadiazine conjugates could serve as potential leads in the development of personalized cancer therapeutics.

**Summary:** The present study describes the exploration of small molecules based on heterocyclic scaffolds for tubulin target based development of anticancer agents.

## Introduction

Cancer, the condition resulting from unbridled progression of cell division, is a fatal affliction to human kind causing one death out of 7 total deaths worldwide. The gloomy prediction of high surge in new cancer cases to 21.7 million by 2030 further underlines the need of concerted endeavours to contain and cure this disease (Kamal et al., 2015). The seminal discovery of “Salvarsan - Magic bullet” laid the foundation for the present day targeted anticancer chemotherapy (http://www.hematology.org/Thehematologist/Profiles/4513.aspx). Tremendous advances achieved in cancer research in the past few decades have also been accompanied by major challenges such as statistical increase in the development of resistance to chemotherapy, limitations in selectivity and associated side effects of anticancer agents (Tiwary et al., 2015) (https://mlsmr.evotec.com/MLSMR_HomePage/index.html). In the development of novel therapeutic agent for cancer, both empirical and rational aspects of drug design are being pursued (Hoskin and Ramamoorthy., 2008). On global level for the last few decades, different organizations have started and are maintaining repositories of diverse synthetic and natural chemical compound to facilitate in the identification and development of potent and nontoxic drugs for pernicious diseases (Holbeck et al., 2010). Most of these repositories like Developmental Therapeutics Program (DTP) by NCI-USA, Molecular Libraries-Small Molecule Repository by NIH-South San Francisco, Open source drug discovery (OSDD) by CSIR-India are focusing on cancer and other diseases like tuberculosis, malaria etc.(http://www.osdd.net/research-development/open-access-repositories). Similarly, the Chemical Synthesis and Drug Supply Program (CSDSP) by NIMH-Bethesda was started with primary purpose to support the scientific research groups working on neurological diseases by synthesizing, purifying and distributing non-commercial and novel chemical entities to assist in psychopharmacology studies and mental health (https://nimh-repository.rti.org/assets/catalog.pdf) (Liu and Campillos., 2014; Srivastava and Srivastava., 2015).

In the last several years, the utilization of the chemical compounds from these repositories to evaluate a specific or shared biological activities/selectivity of chemical entities is being explored in the quest of new drug development. Numerous reports highlight the different biological activities exhibited by a single class of compounds, suggesting that a compound reported for a specific biological activity may or may not be limited to that particular activity with a possibility of it to bind at more than one target at molecular level. Moreover, biological profiles of many compounds in repositories are available in public domain and some of the compounds could be obtained for research purpose. This provides a remarkable and unprecedented support to researchers working in the area of drug development.^7^

Chemical compounds with heterocyclic scaffolds form the pivot of pharmaceutical drug industry as well as new drug discovery and development programs for various diseases. Different heterocyclic moieties are well known for diverse range of pharmacological properties and one of them is nitrogen containing quinazoline pharmacophore. Alkaloids from natural sources contains basic quinazoline skeletal, which is a benzene ring fused to heteroatom containing pyrimidine ring. Peganine is a first quinazoline alkaloid isolated from plant and indolopyridoquinazolinone alkaloid isolated from the plant *Evodia rutaecarpa* in the early 19^th^ century is extensively used as tradition medicine for various ailments related to gastro intestine, absence of menstruation, and postpartum haemorrhage in china (Son et al., 2015; Zayed et al., 2015; Kamal et al., 2011). Extensive reports are available in literature highlighting wide biological activities of quinazoline based derivatives, which includes antimicrobial, anti-inflammatory, anti-malarial, anti HIV, anti-depressant and anti-cancer properties (Kuroiwa et al., 2015). In recent years, the quinazoline based intermediates are reported to exhibit anti-cancer properties by targeting the mitotic spindle during cell division via inhibition of tubulin polymerization, protein kinase of epidermal growth factor receptor (EGFR) (Wang et al., 2014) Erlotinib and Afatinib, the drugs approved by FDA in 2013 are quinazoline derivatives that target EGFR kinases, thereby preventing the growth of cancer cells (Zahedifard et al., 2015). Benzothiadiazines, resulting from the addition of a sulphur atom on quinazoline skeleton, are also reported to exhibit wide range of pharmacological potency. 1,2,4-Benzothiadiazine-1,1-dioxide derivatives are used as diuretic medication and recently the moiety has been explored for different pharmacological properties such as antibacterial, antiviral, anticancer, inhibition of ATP-sensitive potassium channel inhibitors and allosteric modulation of AMPA receptors (Sak 2012; Manasa et al., 2011; Nerkar et al., 2009).

Exploring heterocylic scaffolds for pharmacological potency in our previous studies, we have synthesized a library of quinazoline and benzothiadiazine derivatives and studied their anti-inflammatory and anti-depressant activities in rats (Reddy et al., 1985, 1987). Of late in our laboratory, we explore and investigate novel chemical compounds with varied heterocyclic scaffolds for arresting unregulated growth in cancer cells and the mechanism of action of these new entities (Kamal et al., 2007). The recent advances in the library screening and exploitation of these core moieties have encouraged us to further investigate the quinazoline and benzothiadiazine derivatives stored in our in-house chemical repository for their anticancer activity. For this purpose, we have screened over sixty derivatives for antiproliferative activity against selected human cancer cells to identify promising derivatives. This is followed by mechanistic studies for tubulin polymerization at molecular level for a more focused target based screening for potent anticancer agents.

Microtubule is a dynamic protein made of monomeric α and β tubulin. Many cellular functions are dependent on this microtubule, which includes cell division, cell signalling, cilia and flagellar motions, intracellular transport and intracellular structure etc. Because of its major vital functions in cells it is also called as cytoskeleton and is considered as a desirable target for anticancer agents (Wang and Xie., 2012). In cancer cells, the dynamic stability of tubulin is altered and the tubulin targeting agents act as spindle poison by triggering a mitotic catastrophy resulting in apoptotic cell death. To date several antimitotic compounds are developed which binds to the tubulin at a specific binding site like colchicine, vinca alkaloid, taxol and peloruside binding sites, however most of the compounds bind at colchicine binding site (Wang et al., 2012, 2013).

## Results and discussion

### In vitro antiproliferation activity

From the past few years several research groups are focusing on the quinazoline and benzothiadiazine scaffold to exploit their vast biological activities, especially relating to cancer. This encouraged us to explore the anticancer acitivity of quinazoline (**ATS 1-11**) and benzothiadiazine-1,1-dioxide derivatives (**ATS 12-53**) stored in our in-house chemical repository. Moreover, some of the sulphonamides intermediates (**ATS 54-62**) required in the synthesis of benzothiadiazines were also evaluated. The compounds were carefully selected and were previously reported for their in vivo anti-inflammatory and anti-depression activities. The selected compounds were screened for their anticancer activity on a panel of human cancer cell lines MCF7 and MDA MB-231 (breast carcinoma), DU145 (prostate carcinoma), SKOV3 (ovarian carcinoma) and HepG2 (hepato carcinoma) (ST-1). Among over sixty compounds, four showed potent anti-proliferative activity with GI_50_ values ranging from 0.15 μM to 0.81 μM concentration whereas the other compounds showed moderate activity on the tested cell lines. Interestingly, selectivity towards particular cancer cell line was observed and compounds **ATS 1, 2, & 24** exhibited GI_50_ values 0.15, 0.45, & 0.57, respectively against ovarian cancer SKOV3. Whereas the compounds **ATS-4** and **ATS-7** exhibited GI_50_ of 0.81 and 0.6 μM on DU145 and MCF7, respectively. The GI_50_ values of all the tested compounds are presented in the Table 1. Compound **ATS-7** showed high potency against MCF7 compared to MDA MB-231, in previous literature it was reported that quinazolines exhibited similar activity and many of them are in clinical studies. This encouraged us to select **ATS-7** for further biological studies to understand the mechanism of action.

**Table 1:**
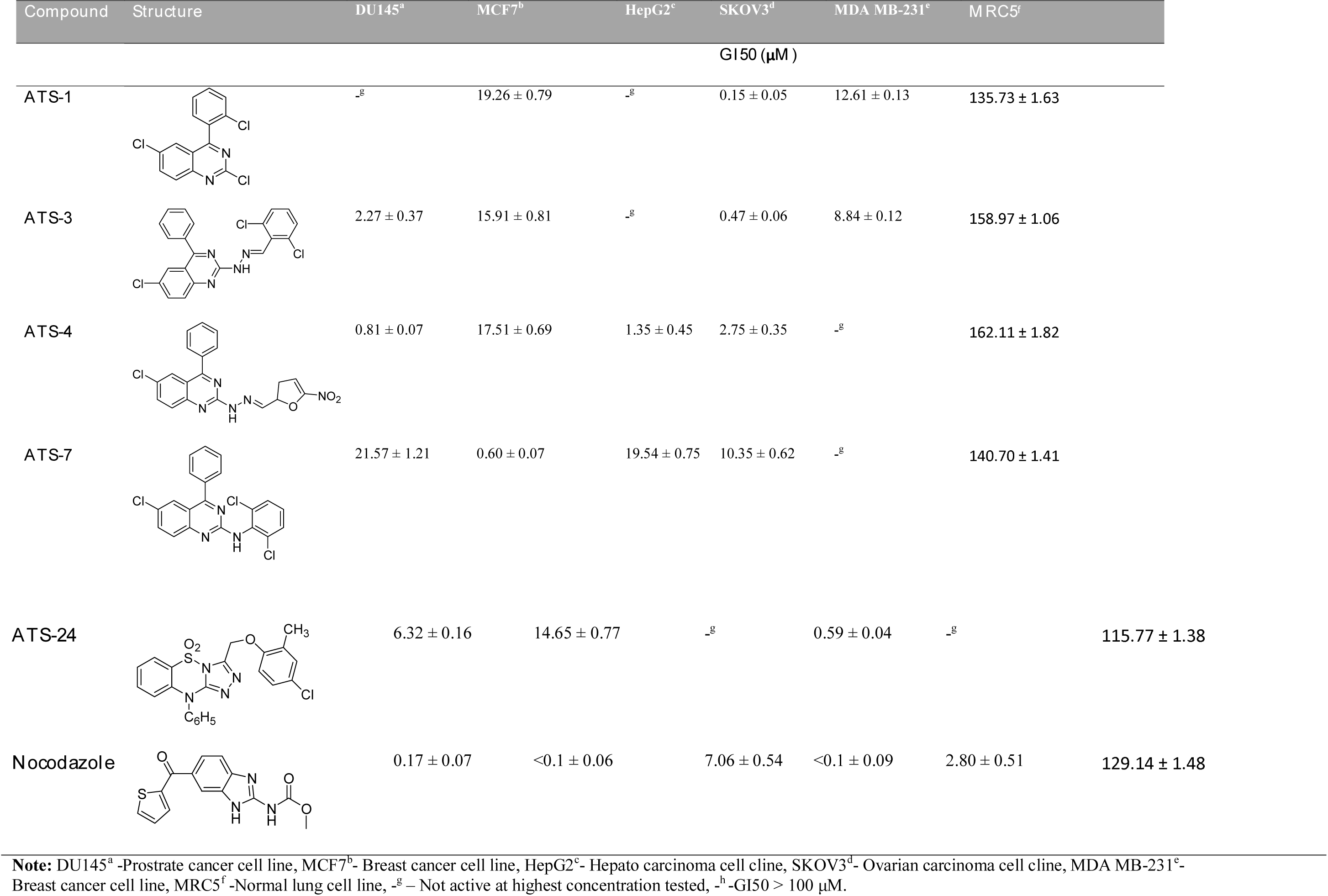
Antiproliferative activity of selected active compounds on cancer and normal cell line.

| Compound | Structure | DU145<sup>a</sup> | MCF7<sup>b</sup> | HepG2<sup>c</sup> | SKOV3<sup>d</sup> | MDA MB-231<sup>e</sup> | MRC5<sup>f</sup> |
| --- | --- | --- | --- | --- | --- | --- | --- |
| **GI50 (μM)** |  |  |  |  |  |  |  |
| **ATS-1** |  | -<sup>g</sup> | 19.26 ± 0.79 | -<sup>g</sup> | 0.15 ± 0.05 | 12.61 ± 0.13 | 135.73 ± 1.63 |
| **ATS-3** |  | 2.27 ± 0.37 | 15.91 ± 0.81 | -<sup>g</sup> | 0.47 ± 0.06 | 8.84 ± 0.12 | 158.97 ± 1.06 |
| **ATS-4** |  | 0.81 ± 0.07 | 17.51 ± 0.69 | 1.35 ± 0.45 | 2.75 ± 0.35 | -<sup>g</sup> | 162.11 ± 1.82 |
| **ATS-7** |  | 21.57 ± 1.21 | 0.60 ± 0.07 | 19.54 ± 0.75 | 10.35 ± 0.62 | -<sup>g</sup> | 140.70 ± 1.41 |
| **ATS-24** | | $6.32 \pm 0.16$ | $14.65 \pm 0.77$ | -<sup>g</sup> | $0.59 \pm 0.04$ | -<sup>g</sup> | $115.77 \pm 1.38$ |
| **Nocodazole** | | $0.17 \pm 0.07$ | $<0.1 \pm 0.06$ | $7.06 \pm 0.54$ | $<0.1 \pm 0.09$ | $2.80 \pm 0.51$ | $129.14 \pm 1.48$ |

**Note:** DU145<sup>a</sup> -Prostrate cancer cell line, MCF7<sup>b</sup> - Breast cancer cell line, HepG2<sup>c</sup> - Hepato carcinoma cell cline, SKOV3<sup>d</sup> - Ovarian carcinoma cell cline, MDA MB-231<sup>e</sup> - Breast cancer cell line, MRC5<sup>f</sup> -Normal lung cell line, -<sup>g</sup> – Not active at highest concentration tested, -<sup>h</sup> -GI50 > 100 μM.

### Effect of compounds on tubulin polymerization

Assay was performed in 384 well black plate contacting 3mg/ml tubulin in the presence of 1 mM GTP and with or without test compound. Fluorescence was measured at 360 nm excitation and 420 nm emission wavelengths. **ATS-7** showed 59% tubulin polymerization inhibition at 3 μM concentration whereas the positive control, nocodazole, showed 67% inhibition at the same concentration. Tubulin IC_50_ was also determined for this molecule (Figure 1).

**Figure 1:**
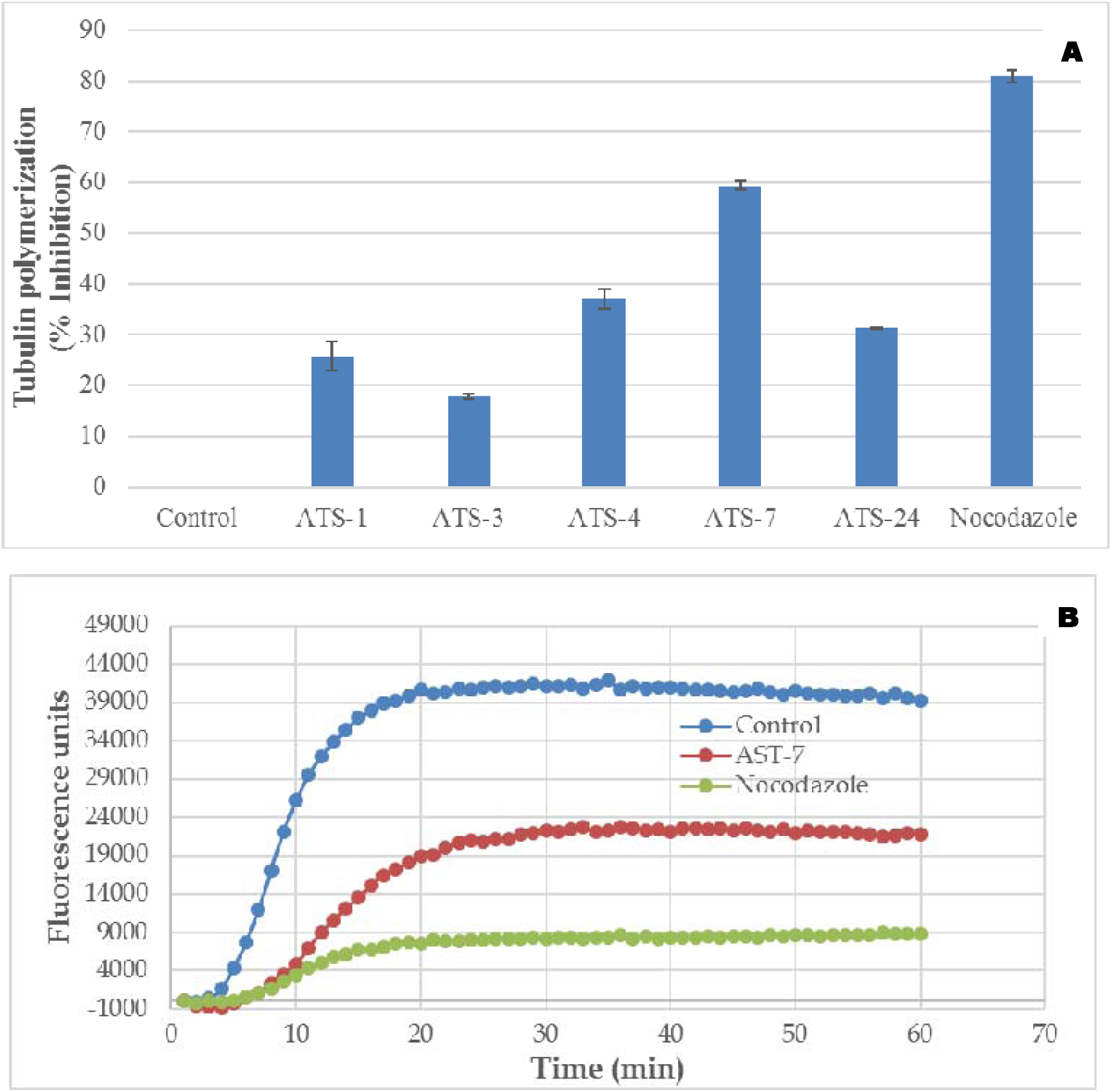
Tubulin polymerization assay: **(a)** Percentage inhibition of tubulin polymerization in presence of drugs (ATS-1, 3, 4,7 &24) at 3 μM concentration was determined by incubating the compound with tubulin in PEM buffer containing 1 mM GTP. Graph was plotted by taking endpoints of the kinetic cycle. Nocodazole was used as a positive control. **(b)** Fluorescence based tubulin polymerization assay of potent compound **AST-7** at 3 μM concentration incubated with tubulin in PEM buffer in presence of 1 mM GTP. Increase in fluorescence was measured at 420 nm emission wavelength when excited at excitation wavelength of 360 nm for a period of 60 mins with one-minute time interval.

### Cell cycle analysis

Most of the anticancer agents arrest the cancer cells at a particular cell cycle phase leading to either apoptosis or necrosis, however some drugs directly induce apoptosis.^9, 10^ Anti proliferative activity and tubulin inhibition studies revealed that compound **ATS-7** showed significant anticancer as well as tubulin inhibition properties. In order to determine whether the effect of **ATS-7** on tubulin polymerization inhibition is translated to similar effect in the cancer cells, cell cycle analysis was performed at two different concentrations in time dependent manner. MCF7 cells were treated with the **ATS-7** and nocodazole at 0.6 and 1.2 μM concentrations, after 12 and 48 h of incubation cells were stained with propidium iodide and analysed for their effect on cell cycle. Clear evidence was observed with the data obtained by cell cycle analysis that **ATS-7** compound arrest the MCF7 cells at G2/M phase. After 12 h of incubation **ATS-7** blocked 16.03%, and 21.24% of MCF7 cells at G2/M phase at 0.6 and 1.2 μM concentration, respectively whereas nocodazole blocked 18.5% and 30.3% cells at G2/M phase under conditions. At 48 h of incubation **ATS-7** and nocodazole showed two-fold enhanced effect by blocking 42.39 %, 48.86%, 22.87%, and 39.73% cells at G2/M phase at 0.6 and 1.2 μM concentrations, respectively (Figure 2). These results demonstrated that **ATS-7** showed comparable and superior activity than nocodazole with the increase in exposure time on cancer cells. It also demonstrated that **ATS-7** anticancer activity of **ATS-7** is due to inhibition of tubulin polymerization and arresting of the cells at G2/M phase as a time and dose dependent manner.

**Figure 2:**
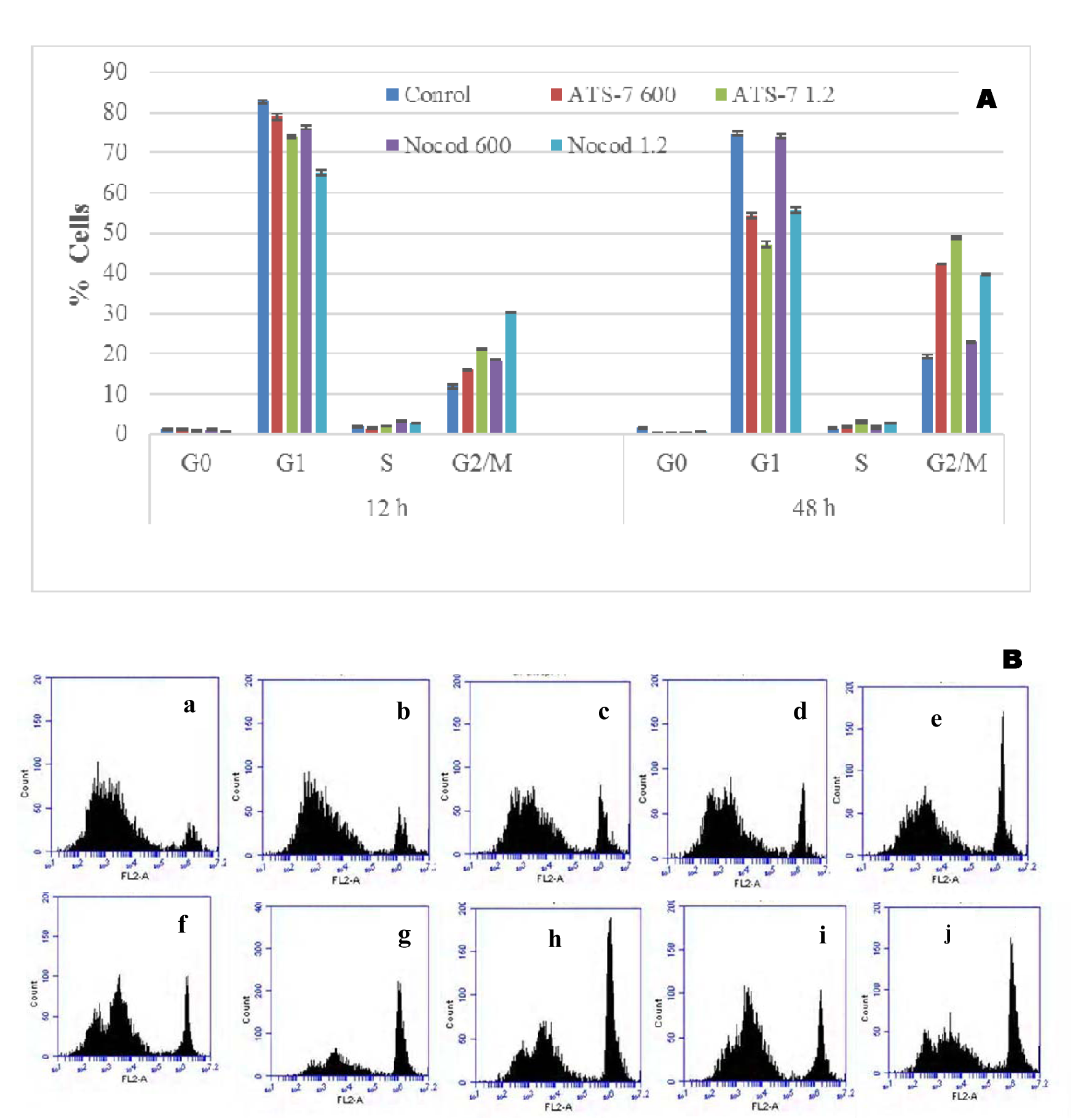
Anti mitotic effect of compound ATS-7: **(A)** Flow cytometric analysis in MCF-7 cell line after treatment with different concentration of test compounds. Percentage of cells were arrested in different phases (G0, G1, S, and G_2_/M) for 12h and 48 h. **(B)** Histograms of the cell cycle arrest at 12 h (A-control cells; B,C-AST-7; D, E- Nocodazole) at 48 h (Fcontrol cells; G, H -**AST-7**; I, J - Nocodazole).

### Immunohistochemistry

Compound **ATS-7** with in vitro tubulin inhibition and cell cycle arrest properties, was expected to disrupt microtubules in cells. To demonstrate cellular microtubule disruption, MCF7 cells were grown on coverslips and treated with **ATS-7** and nocodazole at 0.6 and 1.2 μM concentration. After 48 h of incubation, cells were fixed and treated with primary anti tubulin and rhodamine conjugated secondary antibodies and DAPI for DNA staining as described in experimental section. These cover slips were mounted on slides and analysed under confocal microscope to photograph the tubulin network in cells. Cells treated with **ATS-7** showed altered microtubule network compare to control cells, where well organized microtubule was observed. In both the concentrations **ATS-7** and nocodazole both showed disorganized microtubule network at centre of the cells with dense peripheral microtubules (Figure 3). This demonstrated that **ATS-7** has antimitotic activity comparable with the positive control nocodazole.

**Figure 3:**
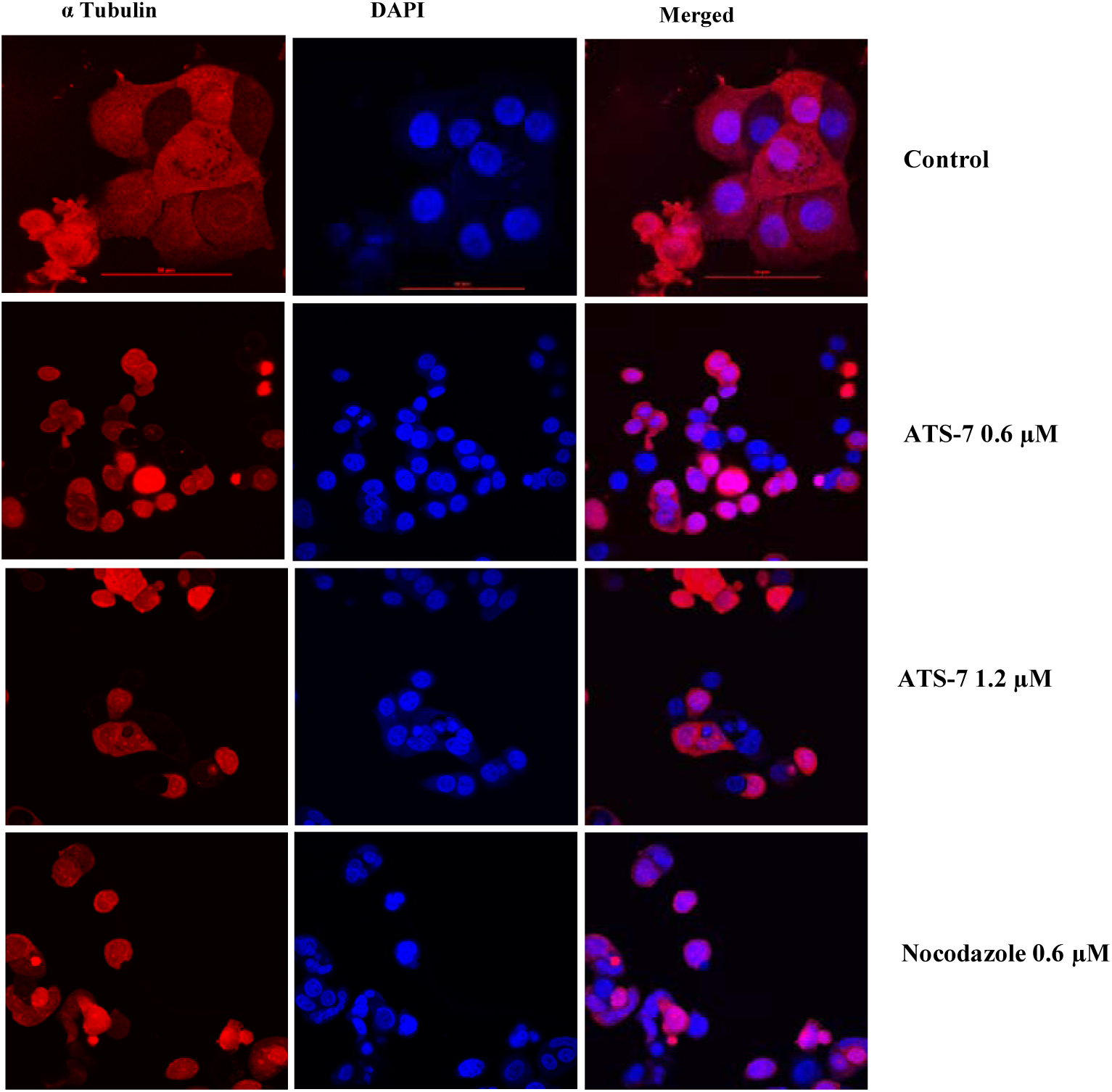
Confocal imaging of cellular microtubules: Immunocytochemistry analysis of the **AST-7** in MCF-7 cell line. The control cells showed normal distribution of tubulin polymerization, whereas the cells were incubated with **AST-7** exhibited disrupted microtubule assembly. DAPI was used as a counterstain for the visualization of the nucleus.

### Soluble versus polymerized tubulin analysis by westernblotting

Inhibition of tubulin polymerization and arrest of cells at G2/M phase in cell cycle analysis was encouraged to analyse the levels of soluble versus polymerized forms of tubulin in MCF7. After incubation of MCF7 cells with compound **ATS-7** (0.6 μM and 1.2 μM) for 48 h, cells were washed with PBS and soluble (free tubulin) and insoluble (polymerized form of tubulin as microtubule) fractions were isolated as described in materials and methods section. Nocodazole and taxol (1 μM) was used as tubulin destabilising and stabilising agents (positive and negative controls respectively). Immunoblot was performed with the isolated fractions and results indicate that cells treated with **ATS-7** and nocodazole showed increased levels of tubulin in soluble fraction compared to the DMSO control, where as in cells treated with taxol showed increased levels of tubulin polymerised fraction (insoluble fraction) (Figure 4). Results suggest that the compound **ATS-7** was a microtubule destabilizing agent.

**Figure 4:**
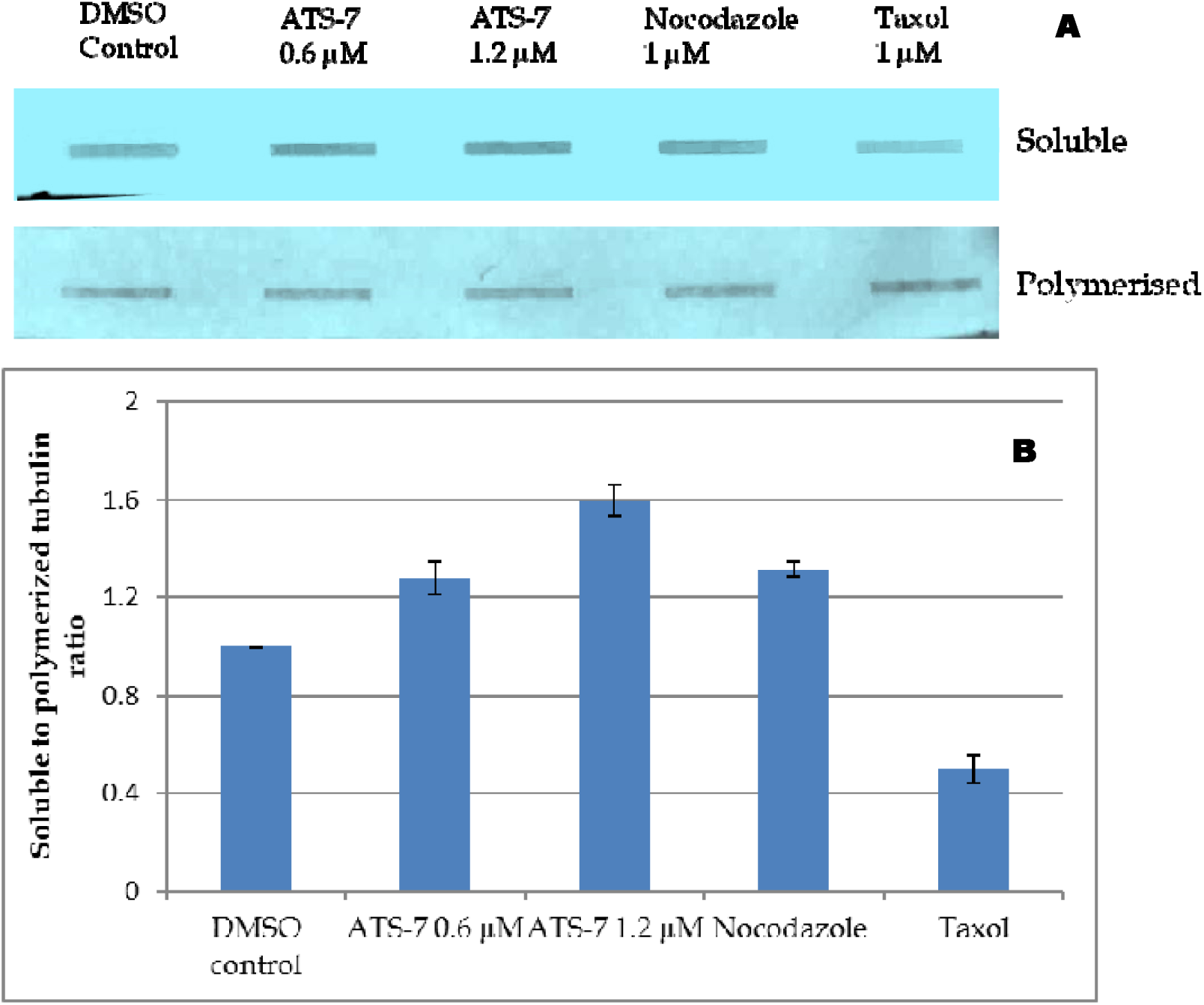
Estimation of cellular levels of tubulin: Effect of **ATS-7** compound on cellular levels of tubulin. **(A)** Western blot analysis soluble and polymerized fraction of MCF7 cells treated with **ATS-7** (0.6 and 1.2 μM). Where Nocodazole and taxol (1 μM) were used as a positive and negative control. **(B)** Densitometry analysis of soluble and polymerized tubulin bands were performed and ration of soluble to polymerized tubulin was represented.

### Mitochondrial membrane potential

Mitochondria is considered as “power house of cells” but studies on mitochondrial function in cells revealed that they are involved in many cellular activities such as lipid modification, calcium level maintenance and cellular death signalling etc. Change in mitochondrial membrane potential (ΔψM) because of any type of cellular stress leads to the activation of mitochondrial mediated apoptosis. This can be measured by using JC-1 dye (cytofluorimetric, lipophilic cationic dye, 5, 5’, 6, 6’-tetraethylbenzimi-diazolylcarbocyanine iodide) which aggregates when cells are healthy where as in cells with altered MMP it is in monomeric form. After incubating the MCF7 cells with **ATS-7**, nocodazole and doxorubicin (0.6 and 1.2 μM concentration) for 48 h, cells were stained with JC-1 dye. The cells were analysed in tecan multimode reader as mentioned in experimental section. Cells treated with **ATS-7** showed 40% decreased signal compared to the healthy control cells (Figure 5). It was evident that **ATS-7** alters the MMP in the MCF7 cells.

**Figure 5:**
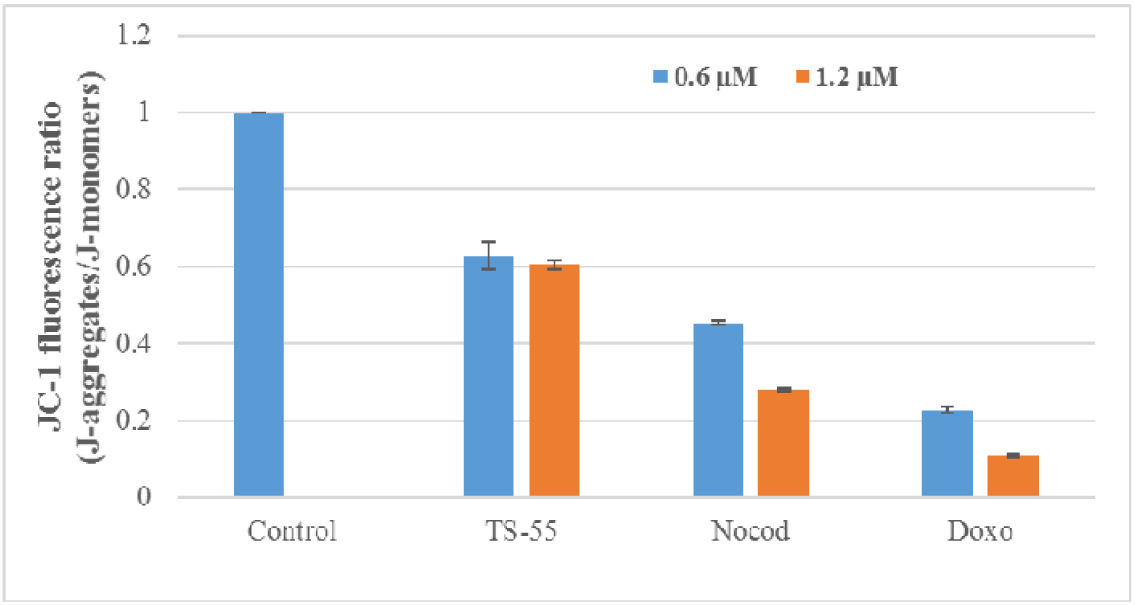
Effect of compound ATS-7 on MMP levels: The loss of mitochondrial membrane potential in MCF-7 cell line treated with different concentrations of **AST-7**, **AST-7/N**, and **AST-7/D**. The untreated sample served as negative control and nocodazole and doxorubicin were used as positive controls. All the experiments were performed in triplicates and the results were expressed as mean ± S.D (n = 3).

### Effect of compound on Caspase 9 activation

Caspases are cysteine-aspartic proteases, activated during abnormal cell functions due to several factors such as radiation, cellular stress, sudden lethal mutations in genomic DNA and chemotherapy leading to apoptosis (Irene et al, 2009). Executioner caspases (caspase 3, 6 and 7) are present in the cells and are activated by the initiator caspases like caspase 2, 8, 9, & 10. Executioner caspases are activated by both caspase 9 and caspase 8 which mediates intrinsic and extrinsic apoptotic pathways, respectively. MCF7 has a defective caspase-3, and caspase 9 plays a crucial role in drug induced apoptosis by activating the other executioner caspases. Therefore, MCF7 cells were treated with **ATS-7** at 0.6 and 1.2 μM concentrations where as positive control nocodazole and doxorubicin were used at 0.6 μM for 48 h. The cells were harvested and analysed for the release of fluorophore from the AFC conjugated LEHD substrate by the enzyme present in the cell lysate. **ATS-7** showed significant caspase 9 activation compared to the positive controls as shown in the Figure 6, which indicated that this compound **ATS-7** induces the mitochondrial mediated apoptosis in MCF7 cells (Figure 6).

**Figure 6:**
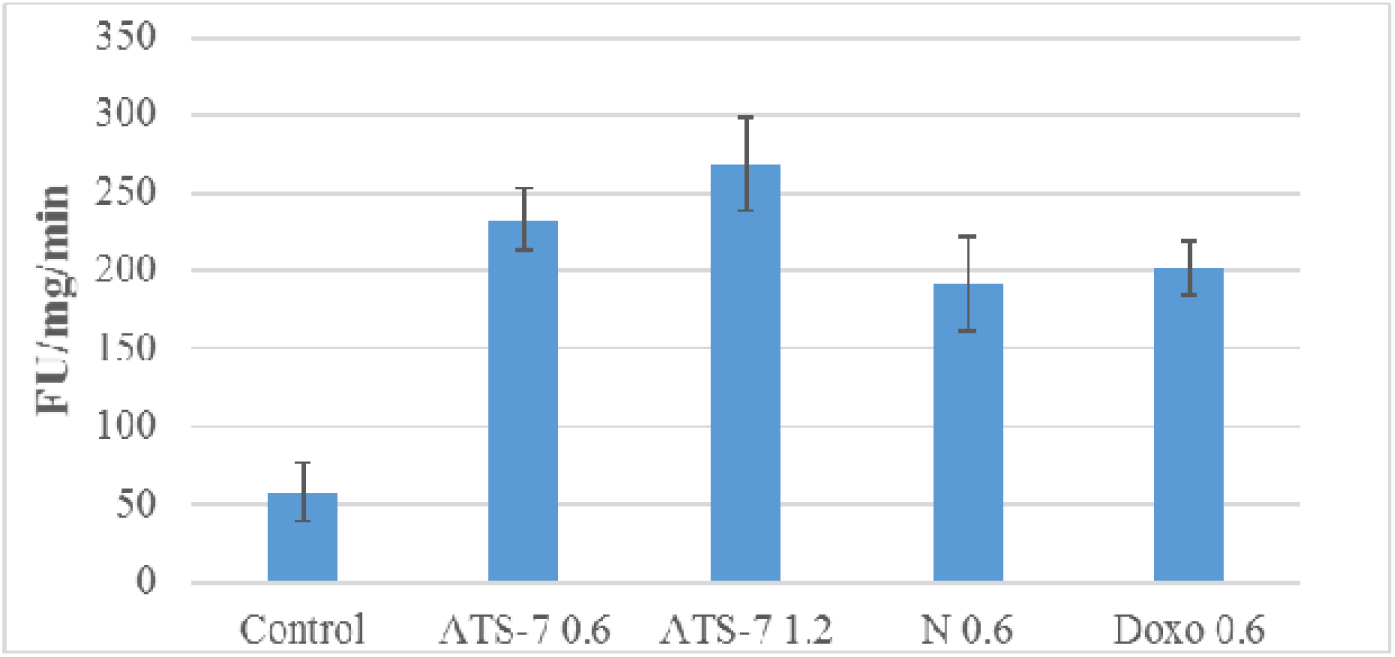
Apoptosis induction in MCF-7 cells treated with ATS-7: The effect of **AST-7** on Caspase −9 in MCF-7 cell lines. The enzymatic activity was assessed in AFC conjugated LEHD substrate. Nocodazole and doxorubicin were used as positive controls. All the experiments were performed in triplicates and the results were expressed as mean ± S.D (n = 3).

### Cooperative index of combination drug treatment

In this investigation combination treatment was studied and MCF7 cells were treated either in combination or alone with **ATS-7**, nocodazole and doxorubicin at GI_50_ concentration of test compound and it double concentration to get final concentrations of 0.6 and 1.2 μM. MTT assay was performed after incubating the cells at four different time intervals like 3, 8, 12 and 24 h. Cooperative index is calculated according the formula given in the experimental section and the death percentage of drug combination is represented in figure 7 whereas CI index is presented in table 2. The data demonstrated that at the lower concentration of **ATS-7/nocodazole** and **ATS-7/doxorubicin** combinations up to 12 h incubation showed synergistic effect whereas at 24 h of incubation showed antagonistic effect and at higher concentration showed antagonistic effect.

**Figure 7:**
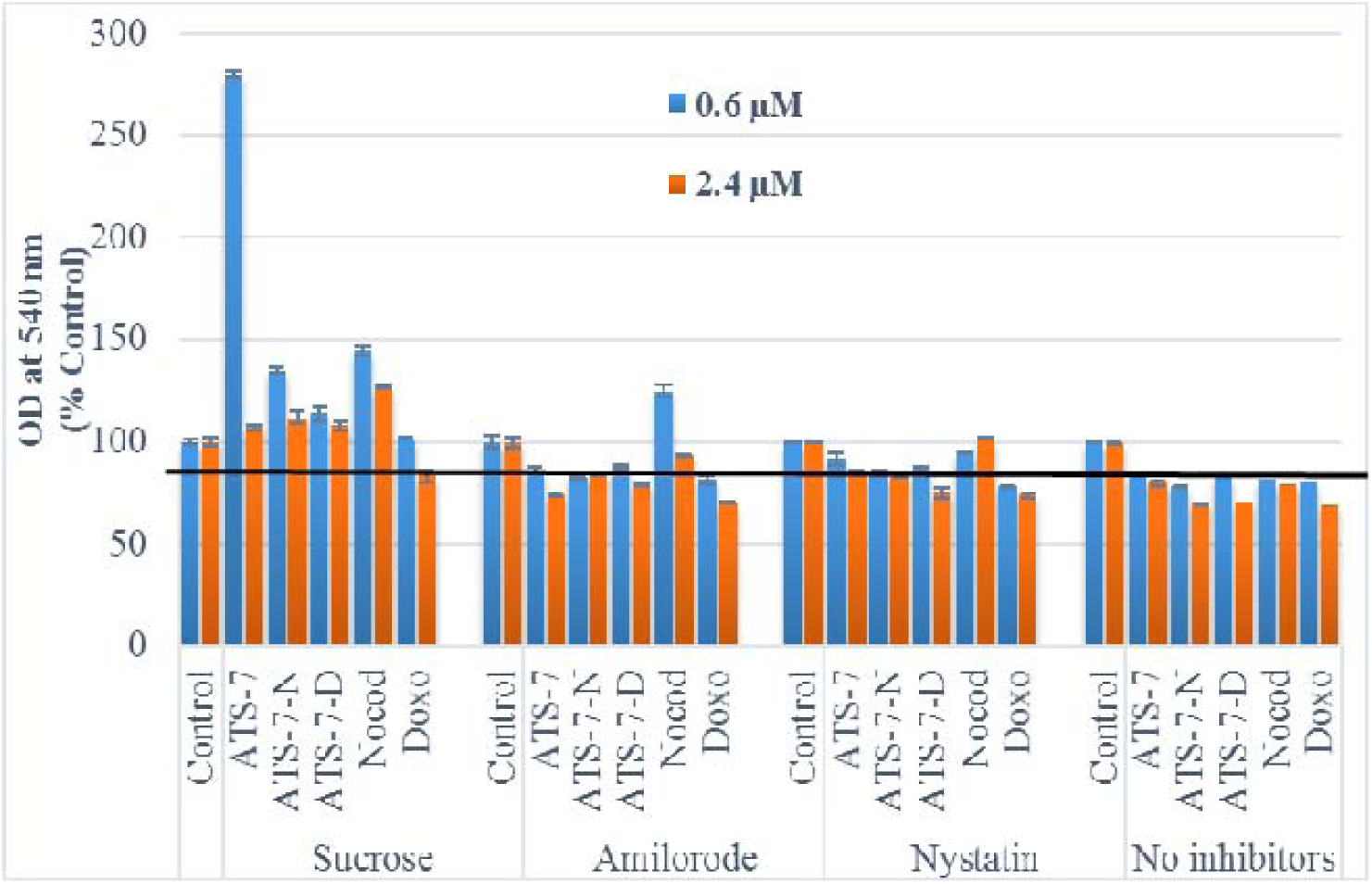
Cellular uptake of ATS-7 by endocytic pathway: **(A)** Endocytic pathway determination for the compound **ATS-7** alone or in combination on MCF-7 cells. MTT assay was performed as described in the experimental section and cell viability was measured at 540 nm. **(B)** Endocytic pathway determination for the compound **ATS-7** alone or in combination on MCF-7 cells. Cellular levels of the drug and the standard was determined by multimode reader at respective wavelengths as described in the experimental section and amount of the drug was calculated from the respective standard graph.

**Table 2:** Cooperative index.

| CI | ATS-7 |  | Incubation time | Effect |
| --- | --- | --- | --- | --- |
| | 0.6 μM | 1.2 μM | | |
| Nocodazole | $0.48 \pm 0.05$ | $1.34 \pm 0.23$ | 3 h | Synergistic |
| | $0.56 \pm 0.02$ | $1.77 \pm 0.15$ | 8 h | Synergistic |
| | $0.95 \pm 0.03$ | $1.81 \pm 0.20$ | 12 h | Synergistic |
| | $2.00 \pm 0.10$ | $1.67 \pm 0.31$ | 24 h | Antagonistic |
| Doxorubicin | $0.13 \pm 0.04$ | $1.62 \pm 0.12$ | 3 h | Synergistic |
| | $0.97 \pm 0.02$ | $2.09 \pm 0.25$ | 8 h | Synergistic |
| | $0.57 \pm 0.03$ | $2.15 \pm 0.22$ | 12 h | Synergistic |
| | $1.82 \pm 0.12$ | $2.22 \pm 0.33$ | 24 h | Antagonistic |

### Endocytic pathway of ATS-7

Bioavailability of drug by cellular uptake and internalization is crucial for its efficacy. Macromolecules cellular uptake is achieved by various mechanisms and internalized drugs/molecules fate is dependent on its uptake mechanism. Drug transport in to the cells are determined by using endocytic pathway inhibitors.^30^ Generally, liquids are transported in to the cells by pinocytosis and it is further classified as macro pinocytosis, clathrine mediated pinocytosis and caveolae mediated pinocytosis. MCF7 cells were incubated with the inhibitors of three endocytic pathways as described in the experimental section for 1 h and then coincubated with **ATS-7**, nocodazole and doxorubicin individually at two different concentrations i.e. 0.6 and 2.4 μM for 2 h. At the end of the experiment MTT assay was performed and internal percentage drug uptake was determined by using multimode plate reader. It was observed that the **ATS-7** uptake was restricted when clathrin mediated endocytic pathway was inhibited, the positive control doxorubicin may also use the clathrin mediated endocytic pathway as shown in the Figure 7. However, nocodazole uptake was restricted in all the endocytic pathway inhibitors treated wells.

## Discussion

Several literature precedence highlights the anticancer properties of quinazoline derivatives and their advantages over the present clinical agents (Sak 2012; Manasa et al., 2011; Nerkar et al., 2009). In one of the studies, Zahedifard et al reported that the modified quinazolines exhibited significant cytotoxicity against MCF7 compared to MDA MB 231 breast cancer cell lines (Zahedifard et al., 2015). In agreement with this, in the present investigation the quinazoline derivatives also exhibited better activity in MCF7 compared to MDA MB-231. Compound **ATS-7** showed high potency against MCF7 with little activity against MDA MB-231 cell line.

Several reports have been found that quinazoline based anti-inflammatory, anti-depressant and VDA’s showed potent anti-tubulin activity. Recently Kenta Kuroiwa et al, Xiao-Feng Wang et al and Virginia Spanò et al. reported anti mitotic activity of quinazoline scaffold molecules (Wang et al., 2012, 2012, 2013, 2013) which encouraged us to investigate the active compounds for their in vitro tubulin polymerization inhibition property (Kuroiwa et al., 2015; Wang et al., 2014, 2013, 2013). As previous reports these quinazolin based derivatives are also showed anti mitotic activity. Inhibition of cell cycle progression and apoptosis are the most common causes of cell growth inhibition. Cell cycle progression is induced by various cell cycle proteins such as CDKs and cyclins as they are the key regulators of cell cycle (Park and Lee 2013). Previous studies reported that treatment of quinazoline derivatives arrests cancer cells in G2/M phase (Cubedo et al., 2006). In the present study, the exposure of MCF7 cells to **ATS-7** compound induced cell cycle arrest in G2/M phase and also induces mitotic arrest. Furthermore, the western blot analysis and immuno cycochemistry studies clearly indicated that **ATS-7** alters the spatial arrangement of tubulin and increased the amount of soluble tubulin in MCF7 cells. Some of the quinazoline derivatives induce cellular apoptosis by altering the MMP, which activates the release of cytochrome C from mitochondria to cytosol. The cytosolic cyt C subsequently activates the levels caspase-9 which leads to apoptosis (Nerkar et al., 2009). In addition, several studies (Iyer et al., 1998; Weir et al., 2007) reported that cell cycle arrest at G2/M phase takes place by the induction of cellular apoptosis. Considering this fact, we further investigated the effect of **ATS-7** compound on the levels of MMP and caspase-9 in MCF7 cells. In general, caspases (cysteine aspartase enzyme), a group of intracellular proteases, are responsible for the induction of apoptosis. In the present investigation, the compound **ATS-7** reduced the levels of MMP and increased the levels of caspase-9 in cancer cells which indicated that these derivatives have the capacity to induce apoptosis in MCF7 cells.

Presently, most of the chemotherapeutic treatments are administered in two or three drug combinations for effective chemotherapy. Several in vitro and clinical investigations on the combination drug treatment suggested that various molecular targets are involved in cancer treatment. Riva and co-workers employed a combination treatment of anti-mitotic compound paclitaxel and VPA (valproic acid) on GSC (Glio stem cells) and observed the synergistic effect to develop a useful combination treatment (Riva et al., 2014). In a similar manner, Konecny and workers also performed the combination drug treatment of paclitaxel with other agents such as epirubicin, carboplatin, gemcitabine and vinorelbine separately on MCF7 cells at lower drug concentrations and observed various effects like synergistic, additive and antagonistic of different combinations (Konecny et al., 2001). It was observed that at higher concentrations the effect of combinations was changed. Effect of **ATS-7** compound in combination with nocodazole and doxorubicin on MCF7 cells was performed. Results demonstrated that **ATS-7-N**, and **ATS-7-D** both showed synergistic effect at lower concentration and incubation time whereas, at lower concentration and higher incubation time (24 h) as well as at higher concentration at all tested incubation time antagonistic effect was observed on MCF7 cells.

Cell membrane is the important barrier which allow entry to various intra and extracellular molecules, further it controls the cell signalling and fate of the internalized molecules (Khalil et al., 2006; Haucke 2015). Previously researchers encountered various challenges during high throughput screening of compound libraries in the early steps of drug discovery. Many molecules fail in clinical phase trials because of unavailability of molecule internalization after cellular uptake and accumulation as well as on how the cells respond to its entry (McShane et al., 2015). Many studies revealed that failed or reduced cellular uptake and accumulation of drug is one of the reasons for drug resistance. Romero and co-workers studied the cisplatin and chloride complex cellular uptake mechanism to understand the cellular influx and efflux equilibrium of these anticancer agents in a time dependent manner. Further, they have also discussed the caveolae mediated endocytosis of these anticancer compounds (Romero et al., 2012).^33^ In this connection in the present investigation quinazoline derivative **ATS-7** cellular uptake is observed by clathrin mediated endocytosis whereas standard nocodazole cellular uptake is mediated by all three endocytic pathways.

## Experimental methods

### In vitro anti-proliferative activity

Anticancer properties of the selected compounds from in house library were evaluated by a standard sulpha rhodamine B assay (SRB) on a panel of selected human epithelial cancer cell lines such as MCF7 (Breast carcinoma), DU145 (Prostate carcinoma), SKOV3 (Ovarian carcinoma), HepG2 (Hepato carcinoma), and MDA MB-231 (Breast carcinoma). The assay was performed in 96 well plate and the cells were seeded with a density of 3.5 × 10^4^ in 100 μL in each well in Dulbecco’s modified Eagle’s medium supplemented with 10% foetal bovine serum, 1% antibiotic solution (PSK) in cell culture incubator at 37 °C temperature supplied with 5% CO_2_ and 100% relative humidity. After 24 h of incubation, the cells were treated with the experimental compounds and positive controls nocodazole and doxorubicin of 2 μl diluted in DMSO at different concentrations (5 × 10^-8^, 1 × 10^-7^, 1 × 10^-6^, 5 × 10^-6^, & 1 × 10^-5^ M). Drug exposure was continued for 48 h in CO_2_ incubator, at the end of incubation cells were fixed with 50 μl of 10% TCA (w/v) at 4 °C for 1 h and followed by washing the cells with water for three times to remove the TCA and then 100 μl of 0.4% SRB (w/v) prepared in 1% acetic acid was added in each well and further incubated for 30 min at RT. The plates were washed with 1% acetic acid for three to five times to remove excess and unbound dye from the wells and then plates were air dried. SRB dye bound to the protein was dissolved by 10 mM Trizma base, optical density of the dissolved dye was measured at 565 nm in a multimode plate reader (Tecan multimode reader). The experiment was carried out in triplicates. Increased absorbance of the SRB dye is relative to the percentage growth of the cells and 50% growth inhibition concentration of the compounds were calculated by the formula described in NCI protocol, GI_50_= (T-T0/C-T0)*100 where T is the absorbance of treated wells with test compounds at 48 h, T0 is the absorbance of wells at time zero and C is the absorbance of the control wells at 48 h (Ashraf et al., 2016; Kamal et al., 2010).

### Tubulin polymerization assay

BK011P kit from Cytoskeleton, Inc. was used to perform fluorescent based tubulin polymerization inhibition assay. According to the manufacturer’s protocol inhibitor detection reaction mixture was prepared for a total volume of 10 μl containing 3 mg/ml porcine brain tubulin, 1 mM GTP and a fluorescence reported in PEM buffer at pH −6.9 (80 mM PIPES, 0.5 mM EGTA, 2 mM MgCl2). In the presence and absence of test compounds at a concentration of 3 μM the increase in fluorescence of reaction mixture was measured at 37 °C at 420 nm emission wavelength by exciting at 360 nm wavelength for every 1 min till 60 min in multimode plate reader. Nocodazole was used as a positive control. Increase in fluorescence was due to the incorporation fluorophore in to the microtubules polymerization.

Tubulin IC_50_ was also determined in the same experimental condition as mentioned above. The concentration of compound where 50% of tubulin polymerization was inhibited compared to the control is known as tubulin IC_50_. Various concentrations like 0.5, 1, 3, 6, and 9 μM of compound **ATS-7** was added in the reaction mixture. Increase in fluorescence as the polymerization progresses was monitored by tecan multimode plate reader at 37 °C for 1 h at 1 min time interval at 420 nm emission wavelength when excited at 360 nm wave length. Nocodazole is used as a positive control (Tecan multimode reader) (Kamal et al., 2012; Kamal et al., 2011).

### Cell cycle analysis

Human breast cancer cells MCF7 were seeded in 6 well plates, after 24 h the cells were treated with the test compound **ATS-7** and a standard drug nocodazole at 0.6 & 1.2 μM concentration for 12 h and 48 h. DMSO was used for appropriate controls. At the end of incubation, cells were trypsinized and harvested by centrifugation for 1 to 2 min at 2000 rpm. The pellets were washed with PBS for 3 to 4 times and fixed in ice cold 70% ethanol for 30 min at 4 °C. Ethanol was removed by centrifugation and cells were stained with 1 ml of DNA staining solution containing 10 mg propidium iodide, 0.1 mg trisodium citrate, 0.03 ml triton X 100 in 100 ml of sterile water. After 30 min of incubation in the staining solution the cells were analysed for the DNA content by flow cytometry (Becton Dickinson FACS Caliber instrument). Histograms were analysed with the software provided by the instrument (Ashraf et al., 2016; Kamal et al., 2011).

### Immuno cytochemistry

MCF7 cells at a density of 1 × 10^4^ cells were seeded on the 18 mm sterile coverslips placed in 6 well plates and incubated for 24 h in a CO_2_ incubator. In the presence and absence of the test compound **ATS-7** at 0.6 and 1.2 μM concentrations, cells were incubated for 48 h in a CO_2_ incubator. Cells were incubated with DAPI for 30 min to stain the DNA. At the termination of experiment, cells were fixed with the mixture of paraformaldehyde, glutaraldehyde solution in 1 ml PBS at a concentration of 3% and 0.02%, respectively for 15 min and permeabilized by dipping them in ice cold 100% methanol for 10 sec. The coverslips were blocked by incubated for 1 h with 1% BSA in PBS and later coverslips were incubated with mouse monoclonal primary anti α tubulin antibody followed by rhodamine conjugated secondary anti-mouse IgG antibody. Coverslips were mounted on a glass slide using DPX fmount and analysed using Nikon A1R confocal microscopy, equipped with rhodamine and DAPI settings. Images were taken at a same scale to analyse the microtubule integrity in test compounds compared to control and positive control nocodazole at 0.6 μM.

### Cellular levels of tubulin

MCF7 cells were seeded in 6-well plates at a density of 2 × 10^5^ in DMEM and treated with or without test compfound **ATS-7** at 0.6 & 1.2 μM concentration. The cells were harvested after 48 h of incubation followed by washing with 1X PBS at 24 °C. Further, the cells were lysed with tubulin lysis buffer (0.1 M Pipes, 1 mM EGTA, 1 mM MgSO4, 30% glycerol, 5% DMSO, 5 mM GTP, 0.125% NP-40, and protease inhibitors (Roche Diagnostics GmbH, Mannheim, Germany), including aprotinin [200 units/ml], pH 6.9) for 30 min at 24 °C temperature. Lysates were centrifuged at 15000g at 24 °C for 30 min, supernatant containing soluble tubulin was collected in separate tube and pellet was suspended with equal volume of lysis buffer containing polymerized tubulin. Pellet and soluble fractions were boiled in lamelli sample buffer for 10 min at 95 °C. Samples were resolved by 10% SDS-PAGE, further immunoblotting was performed after transferring the protein bands on to the nitrocellulose membrane with primary anti α tubulin antibody followed by HRP conjugated secondary anti-mouse IgG antibody.anti-body. Finally, the blot was analysed using ECL prime western blotting mixture supplied by GE health care on photographic film. Quantitative analysis of the soluble and polymer fractions was done by densitometry. Nocodazole and taxol were used as a positive and negative control, respectively at 0.6 μM. Amount of total protein in the soluble and pellet fractions were estimated by Broadford reagent from sigma to normalize the protein before loading the samples in SDS/PAGE gel (Ashraf et al., 2016; Kamal et al., 2012).

### Mitochondrial membrane potential

One of the key functions that altered when the cancer cells are treated with anticancer agents is mitochondrial membrane potential. Most of the anticancer compounds that induce apoptosis will affect the integrity and functioning of mitochondrial membrane. MCF7 cells were analysed for the loss of mitochondrial membrane potential after treatment with the test compound **ATS-7** at 0.6 and 1.2 μM concentrations. Assay was performed using JC-1 dye from Cyman chemicals (CAYM15003-1) and cells were incubated with 0.5 ml JC-1 dye at 37°C in a 5% CO_2_ incubator for 15-30 min. Cells were washed with PBS for 3 times and cells were harvested in 200 μl of assay buffer from each well, cell suspension was centrifuged for 5 min at RT at 1200 rpm and resulted pellet was suspended in 200 μl of 1x assay buffer. 100 μl of cell suspension was transferred in to the 96 well black plate and the cells analysed by fluorescence multimode plate reader. Healthy cells form JC-1 aggregates and strong red fluorescence intensity was measured at 595 nm emission wavelength when excited at 535 nm wavelength, unhealthy cells or altered mitochondrial membrane cells JC-1 monomers exists and strong fluorescence was measured at 535 nm emission wavelength when excited at 485 nm wavelength. The ratio J-aggregates to J-monomers directly implies to the mitochondrial membrane potential of the cells (Budihardjo et al., 1999).

### Caspase 9 Assay

As previously reported that breast cancer cells MCF7 does not express caspase 3 but they undergo apoptosis via caspase 9. MCF7 cells were seeded in 6 well plates, after treatment with or without test compounds and positive control nocodazole for 48 h followed by harvesting of cells in PBS by centrifugation. The cells were suspended in 200 μl of ice cold 1X lysis buffer on ice for 10- 20 min. Supernatant was collected by centrifugation at 13,200 rpm for 20 min at 4°C. Caspase 9 expression in all the samples were measured in 96 well black polystyrene plate in 2X assay buffer containing 50μl cell lysate in the presence of 40 μM of caspase substrate. Increase in fluorescence was measured when caspase 9 cleaved the AFC from Ac-LEHD-AFC at emission wavelength of 505 nm when excited at 400 nm for a period of 1 h at an interval of 5 min. Amount of total protein in all the samples were estimated by Broadford reagent from sigma to normalize the protein (Ye et al., 2001; Mc Gee et al., 2002).

### Combination drug treatment

Cooperative index (CI) is defined as the in vitro cytotoxicity determination of combination drug treatment on a human cancer cell lines. This assay was performed to determine the synergy, antagonism or additive effect of the test compound in combination with other drug at a given exposure time to cells. MCF7 cells were seeded in 96 well plate in a complete medium and treated with test compound and other positive controls nocodazole and doxorubicin individually at 0.6 μM (IC_50_ concentration of **ATS-7**) and 1.2 μM (fixed multiple 2 times of the IC_50_ concentration). Combination treatment also performed at the same concentrations mentioned above in 1:1 ratio. MTT assay was performed at various drug exposure times as previously described. Cooperative Index (CI) was calculated as mentioned in the previous literature, sum of cell death percentages obtained for individual drug treatment divided to the percentage of cell death of combination drug treatment (CI = ATS-7 cell death% + positive control cell death%/Combined treatment cell death%). If CI value is greater than 1 then it has an antagonism effect similarly for additive and synergy effect CI value should be less than 1 and equals to 1 respectively (Riva et al., 2014; Hahm et al., 2001).

### Endocytic pathway determination

Basically two major endocytic pathways are known to exist, namely phagocytosis and pinocytosis. Phagocytosis is one in which larger and solid macromolecules/particles are engulfed by phagosome (referred as cell eating) where as in pinocytosis liquid/solvent material is uptaken by cells (referred as cell drinking). Further pinocytosis is classified to two major groups called micropinocytosis (clathrin-mediated and caveolae-mediated endocytosis) and macropinocytosis. The MCF7 cells were seeded in two replica plates of 24 well plates at a density of 1 × 10^5^, the cells were co-incubate with 154 mg/mL of sucrose (inhibitor of clathrin-mediated endocytosis), 54 μg/mL of nystatin (inhibitor of caveolae-mediated endocytosis) and 133 μg/mL of amiloride (inhibitor of macropinocytosis) for 1 h at 37°C followed by test compounds and positive controls doxorubicin alone and in combination with test compound at 0.6 & 1.2 μM concentration for 2 hours. One replica plate was used for MTT assay and the cells in the other replica plate were then rinsed with PBS at 4°C (3 times). The amount of ATS-7 and doxorubicin inside the cells were analysed by multimode plate reader at 340 nm and 480 nm wave lengths respectively (Khalil et al., 2006; Ding et al., 2015).

## Acknowledgements

We thank Prof Andrew D. Miller, Dr. Maya Thanaou, Centelles Miguel, Michael Wright and Isma Ali from King’s College, London for their support and technical assistance in confocal microscopy & other experiments. T.B.S thank UKIERI council and CSIR-IICT.

## Author contributions

T.B.S. designed and performed the experiments. M.S.M and Z.S.S interpreted the experiments and also helped in the manuscript preparation. S.R.R. assisted in SRB and FACS experiments. A.K conceived, designed and supervised the study and all authors reviewed the manuscript.

## Supplementary information

Supplementary information available

## Graphical Abstract

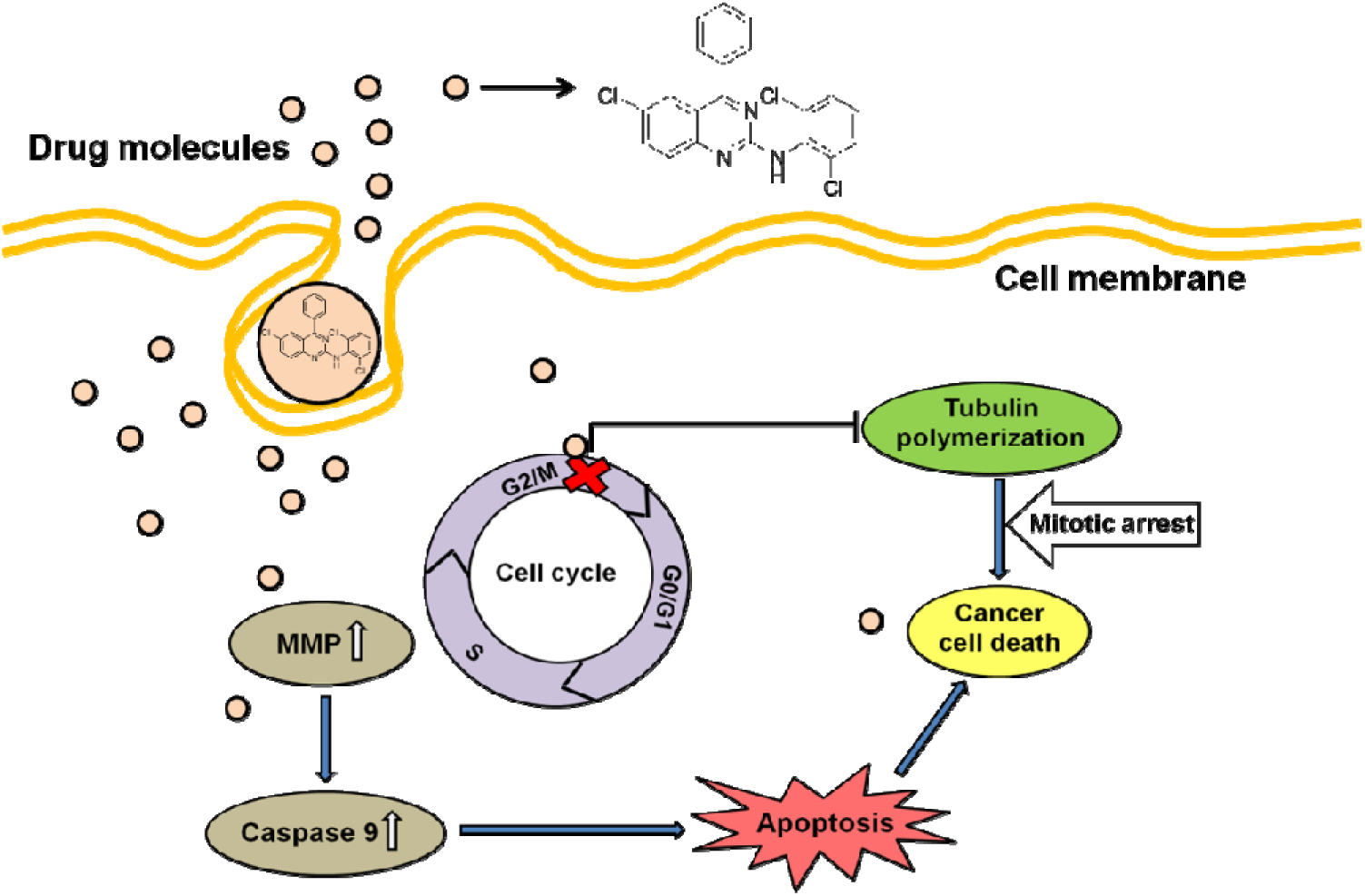

---

**STable 1:** GI_50_ on four different cancer cell lines and values are presented in μM concentraton.

| Compound | DU145<sup>a</sup> | MCF7<sup>b</sup> | HepG2<sup>c</sup> | SKOV3<sup>d</sup> | MDA MB-231<sup>e</sup> |
| --- | --- | --- | --- | --- | --- |
| ATS 1 | -<sup>f</sup> | 19.26 ± 0.79 | -<sup>f</sup> | 0.15 ± 0.05 | 12.61 ± 0.13 |
| ATS 2 | -<sup>f</sup> | 21.68 ± 1.36 | 21.00 ± 0.57 | -<sup>f</sup> | 22.68 ± 1.70 |
| ATS 3 | 2.27 ± 0.37 | 15.91 ± 0.81 | -<sup>f</sup> | 0.47 ± 0.06 | 8.84 ± 0.12 |
| ATS 4 | 0.81 ± 0.07 | 17.51 ± 0.69 | 1.35 ± 0.45 | 2.75 ± 0.35 | -<sup>f</sup> |
| ATS 5 | -<sup>f</sup> | 15.89 ± 0.97 | -<sup>f</sup> | -<sup>f</sup> | -<sup>f</sup> |
| ATS 6 | 6.13 ± 0.49 | 13.83 ± 0.82 | 7.84 ± 0.39 | -<sup>f</sup> | -<sup>f</sup> |
| ATS 7 | 21.57 ± 1.21 | 0.60 ± 0.07 | 19.54 ± 0.75 | 10.35 ± 0.62 | -<sup>f</sup> |
| ATS 8 | 11.02 ± 0.34 | 45.51 ± 1.52 | -<sup>f</sup> | 12.48 ± 0.55 | -<sup>f</sup> |
| ATS 9 | 5.30 ± 0.15 | 36.61 ± 1.36 | 19.96 ± 0.96 | 7.11 ± 0.42 | 18.34 ± 1.09 |
| ATS 10 | 13.32 ± 0.37 | -<sup>f</sup> | -<sup>f</sup> | -<sup>f</sup> | 8.76 ± 0.21 |
| ATS 11 | $8.61 \pm 0.29$ | $10.53 \pm 0.88$ | $_{-f}$ | $15.55 \pm 0.95$ | $_{-f}$ |
| ATS 12 | $42.56 \pm 1.91$ | $9.73 \pm 0.68$ | $_{-f}$ | $_{-f}$ | $20.86 \pm 1.16$ |
| ATS 13 | $6.64 \pm 0.36$ | $17.61 \pm 0.91$ | $_{-f}$ | $4.94 \pm 0.39$ | $_{-f}$ |
| ATS 14 | $19.35 \pm 0.84$ | $_{-f}$ | $_{-f}$ | $5.55 \pm 0.27$ | $8.33 \pm 0.22$ |
| ATS 15 | $_{-f}$ | $_{-f}$ | $_{-f}$ | $7.18 \pm 0.48$ | $_{-f}$ |
| ATS 16 | $20.27 \pm 1.07$ | $_{-f}$ | $_{-f}$ | $7.53 \pm 0.54$ | $_{-f}$ |
| ATS 17 | $3.15 \pm 0.26$ | $28.75 \pm 1.21$ | $21.09 \pm 1.17$ | $11.59 \pm 0.22$ | $_{-f}$ |
| ATS 18 | $13.35 \pm 0.55$ | $16.14 \pm 0.85$ | $_{-f}$ | $_{-f}$ | $3.27 \pm 0.34$ |
| ATS 19 | $9.42 \pm 0.19$ | $_{-f}$ | $10.04 \pm 0.82$ | $37.16 \pm 1.31$ | $2.36 \pm 0.22$ |
| ATS 20 | $11.70 \pm 0.35$ | $35.13 \pm 1.37$ | $_{-f}$ | $_{-f}$ | $17.58 \pm 0.76$ |
| ATS 21 | $7.16 \pm 0.24$ | $31.72 \pm 1.48$ | $_{-f}$ | $_{-f}$ | $21.26 \pm 1.11$ |
| ATS 22 | $17.39 \pm 0.74$ | $_{-f}$ | $137.20 \pm 2.95$ | $_{-f}$ | $_{-f}$ |
| ATS 23 | $10.50 \pm 0.28$ | $_{-f}$ | $_{-g}$ | $28.31 \pm 1.14$ | $_{-f}$ |
| ATS 24 | $6.32 \pm 0.16$ | $14.65 \pm 0.77$ | $_{-g}$ | $0.59 \pm 0.04$ | $_{-f}$ |
| ATS 25 | $_{-f}$ | $_{-f}$ | $_{-f}$ | $_{-f}$ | $19.59 \pm 1.03$ |
| ATS 26 | $_{-f}$ | $_{-f}$ | $_{-f}$ | $_{-f}$ | $_{-f}$ |
| ATS 27 | $4.16 \pm 0.31$ | $34.62 \pm 1.52$ | $_{-f}$ | $_{-f}$ | $_{-f}$ |
| ATS 28 | $_{-f}$ | $_{-f}$ | $_{-f}$ | $_{-f}$ | $_{-f}$ |
| ATS 29 | $_{-f}$ | $_{-f}$ | $_{-g}$ | $8.24 \pm 0.66$ | $35.15 \pm 1.72$ |
| ATS 30 | $_{-f}$ | $7.85 \pm 0.52$ | $4.83 \pm 0.41$ | $11.47 \pm 0.94$ | $22.27 \pm 1.16$ |
| ATS 31 | $_{-f}$ | $_{-f}$ | $_{-f}$ | $_{-f}$ | $_{-f}$ |
| ATS 32 | $_{-f}$ | $14.97 \pm 0.72$ | $_{-f}$ | $_{-f}$ | $_{-f}$ |
| ATS 33 | $13.64 \pm 0.48$ | $_{-g}$ | $11.35 \pm 0.93$ | $_{-f}$ | $28.52 \pm 1.35$ |
| ATS 34 | $4.33 \pm 0.16$ | $124.95 \pm 2.33$ | $2.19 \pm 0.28$ | $_{-f}$ | $10.64 \pm 0.62$ |
| ATS 35 | $_{-f}$ | $_{-f}$ | $14.94 \pm 0.84$ | $_{-f}$ | $14.12 \pm 0.49$ |
| ATS 36 | $12.56 \pm 0.57$ | $28.50 \pm 1.27$ | $_{-f}$ | $_{-f}$ | $11.79 \pm 0.37$ |
| ATS 37 | $19.50 \pm 0.72$ | $_{-f}$ | $_{-f}$ | $_{-f}$ | $31.00 \pm 1.66$ |
| ATS 38 | $_{-f}$ | $14.38 \pm 0.79$ | $6.75 \pm 0.49$ | $_{-f}$ | $_{-f}$ |
| ATS 39 | $19.94 \pm 0.53$ | $19.94 \pm 0.95$ | $15.76 \pm 1.04$ | $48.17 \pm 1.52$ | $_{-f}$ |
| ATS 40 | $10.25 \pm 0.65$ | $5.17 \pm 0.59$ | $_{-f}$ | $12.63 \pm 0.79$ | $24.02 \pm 1.31$ |
| ATS 41 | $31.86 \pm 1.36$ | $11.09 \pm 0.63$ | $_{-f}$ | $10.57 \pm 0.63$ | $11.02 \pm 0.25$ |
| ATS 42 | $25.18 \pm 1.15$ | $12.25 \pm 0.86$ | $13.01 \pm 0.94$ | $10.53 \pm 0.82$ | $_{-f}$ |
| ATS 43 | $_{-f}$ | $_{-f}$ | $_{-f}$ | $_{-f}$ | $16.88 \pm 0.94$ |
| ATS 44 | -<sup>f</sup> | 12.48 ± 0.74 | -<sup>f</sup> | 43.28 ± 1.46 | -<sup>f</sup> |
| ATS 45 | -<sup>f</sup> | -<sup>f</sup> | -<sup>f</sup> | -<sup>f</sup> | 19.82 ± 1.00 |
| ATS 46 | -<sup>f</sup> | 7.64 ± 0.48 | -<sup>f</sup> | -<sup>f</sup> | 16.02 ± .99 |
| ATS 47 | -<sup>f</sup> | -<sup>f</sup> | -<sup>f</sup> | -<sup>f</sup> | 10.10 ± 0.82 |
| ATS 48 | -<sup>f</sup> | -<sup>f</sup> | 10.09 ± 0.62 | -<sup>f</sup> | 9.22 ± 0.14 |
| ATS 49 | -<sup>f</sup> | 6.02 ± 0.39 | 4.92 ± 0.37 | 18.36 ± 0.99 | -<sup>f</sup> |
| ATS 50 | 3.11 ± 0.18 | 11.38 ± 0.69 | 29.72 ± 1.33 | -<sup>f</sup> | 9.94 ± 0.36 |
| ATS 51 | 32.14 ± 1.26 | 41.21 ± 1.41 | -<sup>f</sup> | 19.54 ± 0.85 | 40.69 ± 2.01 |
| ATS 52 | 9.93 ± 0.62 | -<sup>f</sup> | -<sup>f</sup> | -<sup>f</sup> | 7.28 ± 0.62 |
| ATS 53 | -<sup>f</sup> | 24.71 ± 1.03 | -<sup>f</sup> | -<sup>f</sup> | 11.54 ± 0.17 |
| ATS 54 | -<sup>f</sup> | 6.28 ± 0.38 | 35.37 ± 1.39 | 11.19 ± 0.47 | -<sup>f</sup> |
| ATS 55 | -<sup>f</sup> | 9.57 ± 0.41 | 30.84 ± 1.45 | 13.67 ± 0.59 | -<sup>f</sup> |
| ATS 56 | 26.22 ± 1.06 | -<sup>g</sup> | 34.02 ± 1.28 | -<sup>g</sup> | -<sup>f</sup> |
| ATS 57 | 17.84 ± 0.91 | -<sup>g</sup> | -<sup>g</sup> | 20.29 ± 1.10 | -<sup>f</sup> |
| ATS 58 | -<sup>f</sup> | 13.09 ± 0.93 | 18.92 ± 1.02 | 19.46 ± 1.21 | -<sup>f</sup> |
| ATS 59 | -<sup>f</sup> | 5.26 ± 0.27 | -<sup>f</sup> | 22.22 ± 1.35 | -<sup>f</sup> |
| ATS 60 | -<sup>f</sup> | -<sup>g</sup> | -<sup>f</sup> | 12.11 ± 0.77 | -<sup>f</sup> |
| ATS 61 | -<sup>f</sup> | 4.48 ± 0.52 | -<sup>f</sup> | -<sup>f</sup> | -<sup>f</sup> |
| ATS 62 | 4.72 ± 0.35 | 5.09 ± 0.37 | -<sup>f</sup> | 21.11 ± 1.33 | -<sup>f</sup> |
| Nocodazole | 0.17 ± 0.07 | <0.1 ± 0.06 | 7.06 ± 0.54 | <0.1 ± 0.09 | 2.80 ± 0.51 |
| Doxorubicin | <0.1 ± 0.38 | <0.1 ± 0.27 | <0.1 ± 0.48 | 1.17 ± 0.52 | <0.1 ± 0.25 |

**Note:** DU145<sup>a</sup> -Prostrate cancer cell line, MCF7<sup>b</sup> - Breast cancer cell line, HepG2<sup>c</sup> - Hepato carcinoma cell cline, SKOV3<sup>d</sup> - Ovarian carcinoma cell cline, MDA MB-231<sup>e</sup> - Breast cancer cell line, -<sup>f</sup> – Not active at highest concentration tested, -<sup>h</sup> -GI50 > 100 µM.

